# Assessing the Gene-Disease Association of 19 Genes with the RASopathies using the ClinGen Gene Curation Framework

**DOI:** 10.1101/323303

**Authors:** Andrew Grant, Brandon Cushman, Hélène Cavé, Mitchell W. Dillon, Bruce D. Gelb, Karen W. Gripp, Jennifer A. Lee, Heather Mason-Suares, Katherine A. Rauen, Lisa M. Vincent, Martin Zenker

**Affiliations:** Mindich Child Health and Development Institute and the Departments of Pediatrics and Genetic and Genomic Sciences, Icahn School of Medicine at Mount Sinai, New York, NY; Département de Génétique, Hôpital Robert Debré and Institut Universitaire d’Hématologie, Université Paris Diderot, Paris-Sorbonne-Cité, Paris, France; Icahn School of Medicine at Mount Sinai, Molecular Genetic Testing Laboratory, New York, NY; Nemours/Alfred I. duPont Hospital for Children, Wilmington, DE; Greenwood Genetic Center, Greenwood, SC; Laboratory for Molecular Medicine, Partners Healthcare Personalized Medicine, Cambridge, MA; UC Davis Children’s Hospital, Sacramento, CA; GeneDx, Gaithersburg, MD; Institute of Human Genetics, University Hospital Magdeburg, Germany

**Keywords:** RASopathy, ClinGen, gene curation, genetic, genomics

## Abstract

The RASopathies are a complex group of diseases regarding phenotype and genetic etiology. The ClinGen RASopathy Expert Panel assessed published and other publicly available evidence supporting the association of 19 genes with RASopathy conditions. Using the semi-quantitative literature curation method developed by the ClinGen Gene Curation Working Group, evidence for each gene was curated and scored for Noonan syndrome, Costello syndrome, cardiofaciocutaneous (CFC) syndrome, Noonan syndrome with multiple lentigines (NSML), and Noonan-like syndrome with loose anagen hair (NS/LAH).

The curated evidence supporting each gene-disease relationship was then discussed and approved by the ClinGen RASopathy Expert Panel. Each association’s strength was classified as Definitive, Strong, Moderate, Limited, Disputed, or No Evidence. Eleven genes were classified as definitively associated with at least one RASopathy condition. Two genes classified as strong for association with at least one RASopathy condition while one gene was moderate and three were limited. The RAS EP also refuted the association of two genes for a RASopathy condition. Overall, our results provide a greater understanding of the different gene-disease relationships within the RASopathies and can help guide and direct clinicians, patients and researchers who are identifying variants in individuals with a suspected RASopathy

**GRANT NUMBERS:** Research reported in this publication was supported by the National Human Genome Research Institute (NHGRI) under award number U41HG006834. MZ received support from German Federal Ministry of Education and Research (BMBF): NSEuroNet (FKZ 01GM1602A), GeNeRARe (FKZ 01GM1519A).

## Introduction

The RASopathies are a collective group of phenotypically related conditions caused by germline pathogenic variants in genes within the Ras/mitogen-activated protein kinase (Ras/MAPK) signaling pathway. RASopathy conditions, such as Noonan syndrome (NS) and cardio-facio-cutaneous (CFC) syndrome, typically present with multiple phenotypic features including poor growth, cardiac anomalies, ectodermal abnormalities, neurodevelopmental deficits and increased tumor risk (Nava et al., 2007; Pierpont et al., 2014; Rauen, 2013; Romano et al., 2010; Tidyman & Rauen, 2009). Most conditions within the RASopathies have been historically described as clinically distinct syndromes and strong correlations have been noted between each individual syndrome and a mutated gene(s) or even a specific allele within a gene. Despite these correlations, these conditions share a considerable amount of overlapping phenotypic features that can complicate clinical diagnoses. Furthermore, variable expressivity has been described in individuals and families sharing the same genotype (Allanson & Roberts, 1993; Tartaglia et al., 2002). Therefore, the ClinGen RASopathy expert panel sought to evaluate the current evidence for each gene:condition assertion in order to provide a comprehensive review of Ras/MAPK pathway genes and their causality of a RASopathy condition.

The ClinGen gene curation effort provides formal evidence-based classifications for the association of a gene with a given disease. This type of information is extremely important as clinicians often use molecular testing results to confirm a clinical diagnosis. Therefore, knowing the level of association of a gene with disease can facilitate more accurate clinical diagnoses. This information also allows for more accurate clinical utility and sensitivity in diagnostic laboratory test designs for specific clinical indications. For example, this exercise supports the exclusion of testing for genes with insufficient evidence to support an association with a RASopathy while also informing which RASopathy genes *should* be included when testing for specific RASopathy conditions (e.g. a CFC panel). While this list of genes does not include the entirety of genes that have been associated with RASopathies to date (e.g. *CBL, NF1* and *SPRED1* are not included in this exercise), our work does inform 95 gene:disease associations. Having comprehensive and concise panels is crucial for the practice of molecular medicine, and gene curation has proven to be useful in this endeavor (Strande et al., 2017).

For certain gene-disease associations, extracting accurate and clear phenotypic information from the literature can be challenging; however, the phenotypic and genetic heterogeneity within the RASopathies is particularly confounding for establishing clear gene associations with specific RASopathy conditions. Due to variable expression and high clinical overlap of the features, clinicians who diagnose RASopathy patients have often used the molecular diagnosis to support a specific phenotypic diagnosis in their patients. For example, if a patient displays phenotypes that overlap between NS and CFC syndrome, they may use the finding of a *de novo* variant in *MAP2K2* as supporting evidence that the patient has CFC instead of relying on specific clinical features. While this exemplifies the utility of molecular medicine as a tool for diagnosis, it also highlights the bias for certain genes to be traditionally associated to only certain RASopathy phenotypes. However, given that the complexities of the pathway have not been studied in enough detail to rule out that certain *MAP2K2* variants may lead to an NS phenotype, it is useful to note when evidence supports that a patient with a *MAP2K2* variant may have NS and *not* CFC. While a major part of the utility of the gene curation framework lies in acknowledging all potential associations, RASopathy disorders may have distinguishing features that vary either due to the age of clinical presentation and assessment, generalized phenotypic heterogeneity, or even potentially biased ascertainment. Given these phenotypic similarities overall, a patient diagnosed initially with one entity may transition to a different clinical diagnosis over time as different features manifest or clinical diagnoses are refined. Despite this, there are some nuances and notable differences that can be made among these disorders **(see Table 1).** For example, features like hypertrophic cardiomyopathy are observed across all RASopathies, yet are more commonly observed in patients clinically diagnosed with Costello syndrome (CS) or Noonan syndrome with multiple lentigines (NSML). In general, multiple lentigines are very common, but within the context of a RASopathy they are highly correlated to NSML. Certain hematologic or oncologic manifestations, including solid tumors and transitional cell carcinoma, are highly specific for CS **(Table 1).** Through systematic collection of published literature and other case-level evidence from diagnostic and research laboratories, these nuances and differences in phenotypes enable a better understanding of the larger phenotypic spectrum for each gene. The evidence curated by ClinGen can highlight for clinicians which condition(s) have been associated with a gene and therefore which phenotypic features any given patient may be at risk for. This project also underscores the need for better nosology in the RASopathy field to reduce diagnostic discrepancies between physicians.

**Table 1:**
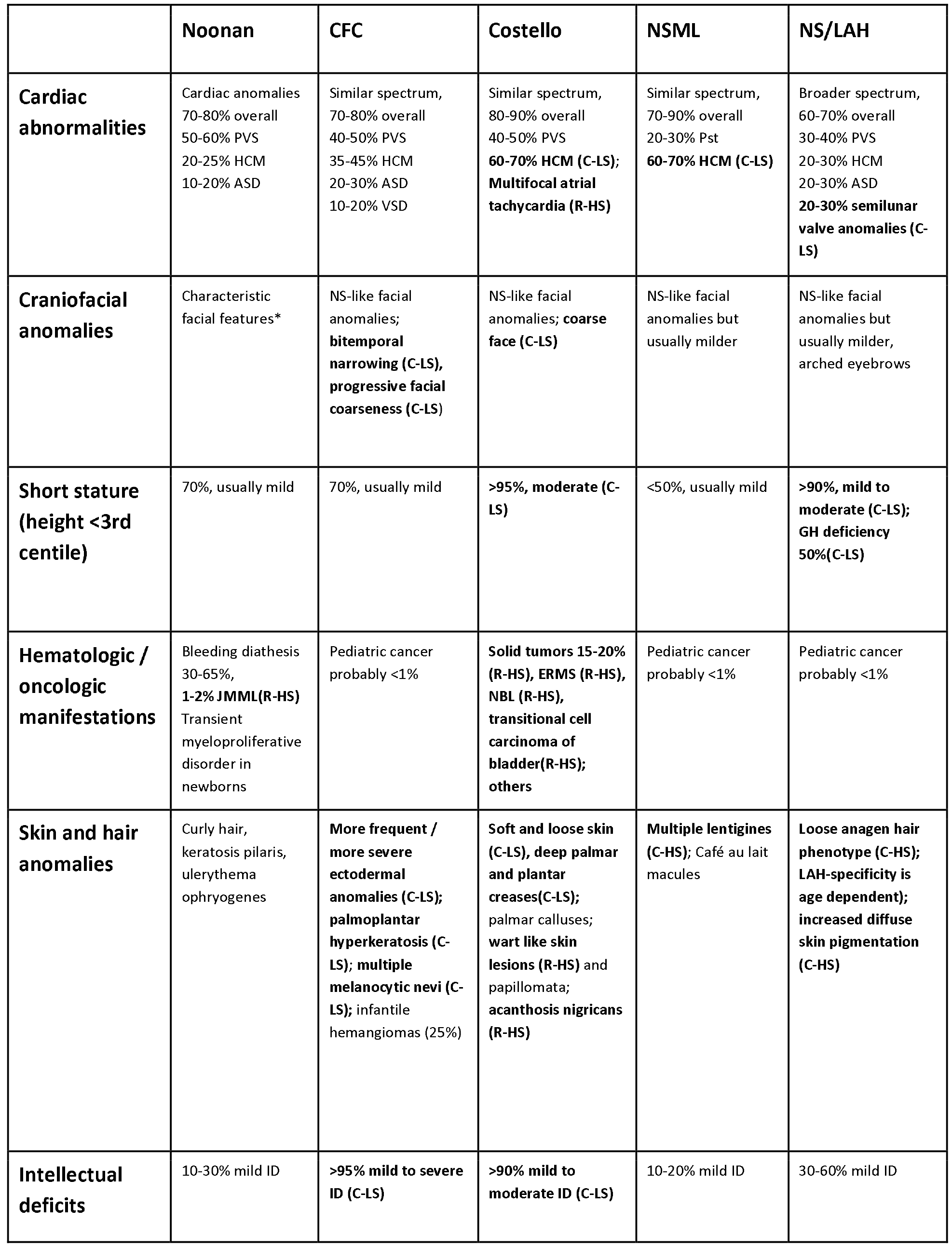
Clinical features of various RASopathy diseases are listed(Allanson & Roberts, 1993; Gelb & Tartaglia, 1993; Gripp & Lin, 1993; Pierpont et al., 2014; Rauen, 1993; Roberts, Allanson, Tartaglia, & Gelb, 2013; Romano et al., 2010). Features that may help to discriminate individual entities from each other are grouped into three categories (bold): 1. Common-Low Specificity (C-LS): relatively common feature (>20%) with low specificity for entity but observed more frequently than in NS; 2. Common-High Specificity (C-HS): relatively common feature (>20%) with high specificity for entity compared to other RASopathies; 3. Rare-High Specificity (R-HS): Rare feature (<20%) with high specificity for entity compared to other RASopathies. ^*^Characteristic facial features of Noonan syndrome include: a broad/ tall forehead, hypertelorism with downslanting palpebral fissures, low-set, posteriorly rotated ears with a thickened helix, a deeply grooved philtrum with high, wide peaks to the vermilion border of the upper lip, a pointed chin, and a short neck with excess nuchal skin or pterygium colli. Facial features of Noonan syndrome change with age. NS, Noonan syndrome; CFC, Cardiofaciocutaneous syndrome; NSML, Noonan syndrome with multiple lentigines; NS-LAH, Noonan syndrome with loose anagen hair; GH, growth hormone; ID, intellectual disability; JMML, juvenile myelomonocytic leukemia; ERMS, embryonal rhabdomyosarcoma; NBL, neuroblastoma.

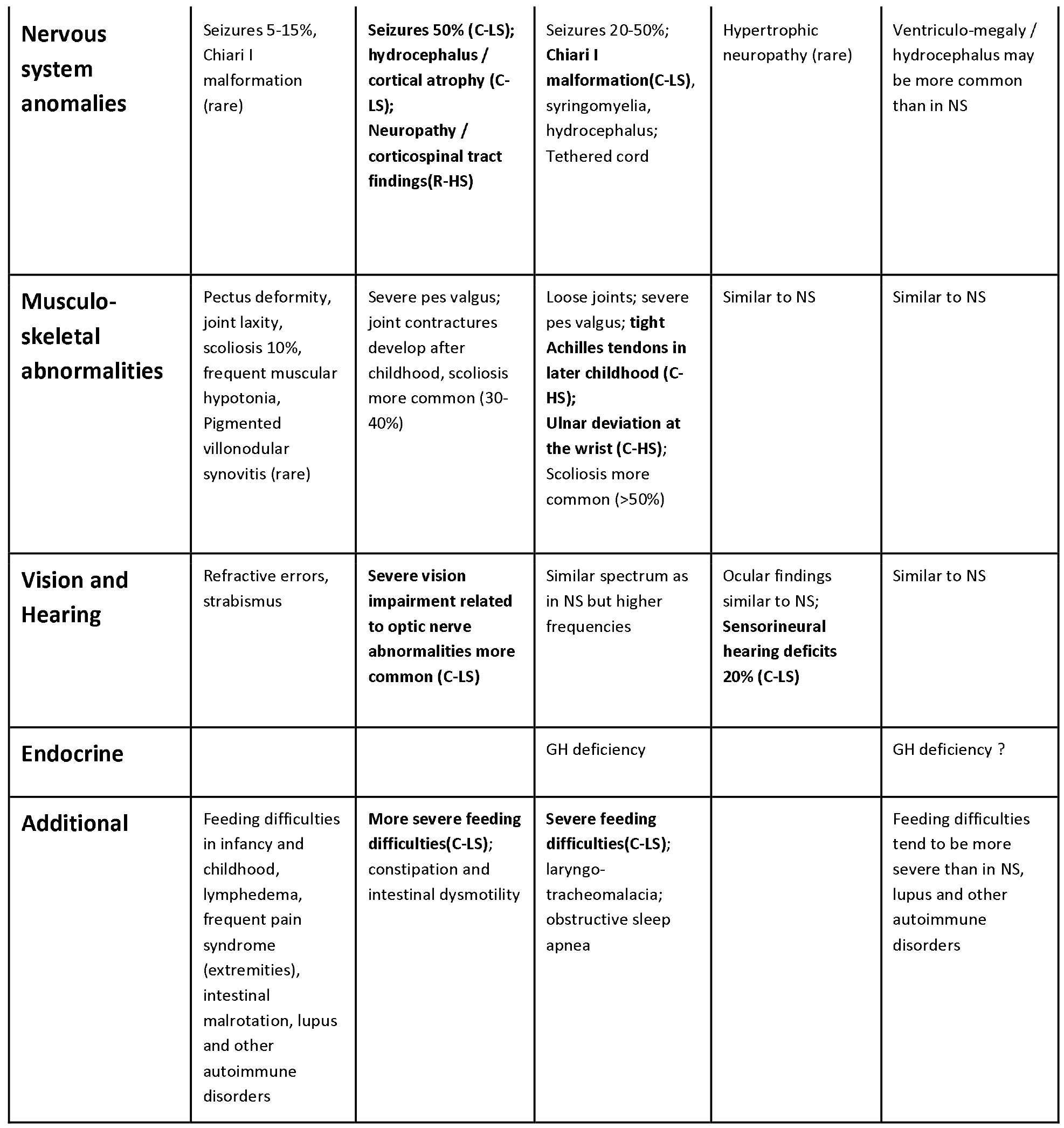

We curated evidence for the association of 19 genes with five specific RASopathy conditions: NS, CFC, CS, NSML, and Noonan-like syndrome with loose anagen hair (NS/LAH). To limit the scope and enable for community feedback of this pilot initiative, our gene list primarily consisted of genes typically associated with a gain of function mechanism leading to a RASopathy in the literature. While many of the gene-disease classifications were curated as definitive and historically linked, we also evaluated genes recently linked to RASopathies, *e.g. PPP1CB* and *LZTR1*, with the aid of additional diagnostic laboratory data publicly available within ClinVar (https://www.ncbi.nlm.nih.gov/clinvar/). We believe that our expert-approved gene-disease validity classifications will be highly informative to patients, researchers, and clinicians interested in the gene-disease associations in the field of RASopathies.

## Methods

The association of each of the 19 genes were classified using the ClinGen Gene Curation Standard Operating Procedures (version 5) to the following specific RASopathy phenotypes: NS, NS/LAH, CFC, CS and NSML (Strande et al., 2017). This framework involves a structured evaluation of published literature to produce a semi-quantitative score relative to the strength of evidence available for a given gene-disease association. In addition, diagnostic laboratory data publicly available within ClinVar (https://www.ncbi.nlm.nih.gov/clinvar/) was also assessed for genes with minimal literature. Genes curated in this study include *A2ML1* (OMIM #610627), *BRAF* (OMIM #164757), *HRAS* (OMIM #190020), *KRAS* (OMIM #190070), *LZTR1* (OMIM #600574), *MAP2K1* (OMIM #176872), *MAP2K2* (OMIM #601263), *MR AS* (OMIM #608435), *NRAS* (OMIM #164790), *PPP1CB* (OMIM #600590), *PTPN11* (OMIM #176876), *RAF1* (OMIM# 164760), *RASA1* (OMIM #139150), *RASA2* (OMIM #601589), *RIT1* (OMIM #609591), *RRAS* (OMIM #165090), *SHOC2* (OMIM #602775), *SOS1* (OMIM #182530), and *SOS2* (OMIM #601247). Each association was classified as Definitive (12-18 points with replication), Strong (12-18 points), Moderate (7-11 points), Limited (0.1-6 points), No Evidence, Disputed or Refuted per the ClinGen Gene Curation criteria (Strande 2017).

After primary curation by a ClinGen biocurator, the evidence for each association was presented to clinical RASopathy experts from several different institutions for review. The evidence and the current clinical validity classification and interpretation supporting the gene-disease or gene-phenotype relationship were discussed at length followed by a vote. If the vote for the classification was unanimous, the association was approved. If the proposed association was unclear or contested, then the association was discussed at length and voting continued until an 80% quorum from all ClinGen RASopathy Expert Panel (RAS EP) members was achieved. In several cases, the RAS EP approved caveat language for describing and refining the term asserting the clinical validity of an association **(Supplementary Table S1).** For example, some Limited associations (*i.e., MAP2K2*:NS-only one scored published case) were predicted by experts to be eventually disputed/refuted, and the group stressed that viewers of those marked “Limited” associations should exercise caution.

## Results

A total of 19 genes were curated for their association with five specific RASopathy conditions. The genes found to be Definitive for at least one RASopathy condition were *BRAF, HRAS, KRAS, MAP2K1, MAP2K2, NRAS, PTPN11, RAF1, RIT1, SHOC2*, and *SOS1*. Clinical phenotypes of cases in the literature or ClinVar were reviewed for consistency with the clinical diagnoses provided at the time of the report. Most genetic evidence scored was that of *de novo* variant occurrences. There were no published case-control studies to score. All genes historically known to participate within the Ras/MAPK pathway or have homologous function to known genes achieved a 0.5 points in the functional score based on the well-established biochemical function and protein interactions of the pathway. Due to the ubiquitous expression of these genes, tissue-specific expression did not provide functional evidence to associate these genes with the RASopathies. Strength of evidence associated with animal models and rescue models was typically downgraded to 0.5 points due to the lack of phenotypic specificity in the models for the discrete RASopathy conditions. Functional alterations from observed variants in patient and nonpatient derived cells were assessed stringently for supporting disease causation and were only scored separately when a distinct assay directly supported a unique mechanism or endpoint for the specific condition being assessed. For example, NSML caused by variation in *PTPN11* results in a predicted neomorphic allele that has reduced catalytic activity (but is not equivalent to a null allele, which is known to cause metachondromatosis) compared to the standard increased activity associated with PTPA/ll-related Noonan syndrome variants (Fragale, Tartaglia, Wu, & Gelb, 2004; Kontaridis, Swanson, David, Barford, & Neel, 2006; Noda, Takahashi, Hayashi, Tanuma, & Hatakeyama, 2016; Oishi et al., 2006; Oishi et al., 2009; Yu et al., 2014). Table 2 summarizes the classifications of all curations along with their semi-quantitative score for specific RASopathy conditions (maximum out of 18).

**Table 2:**
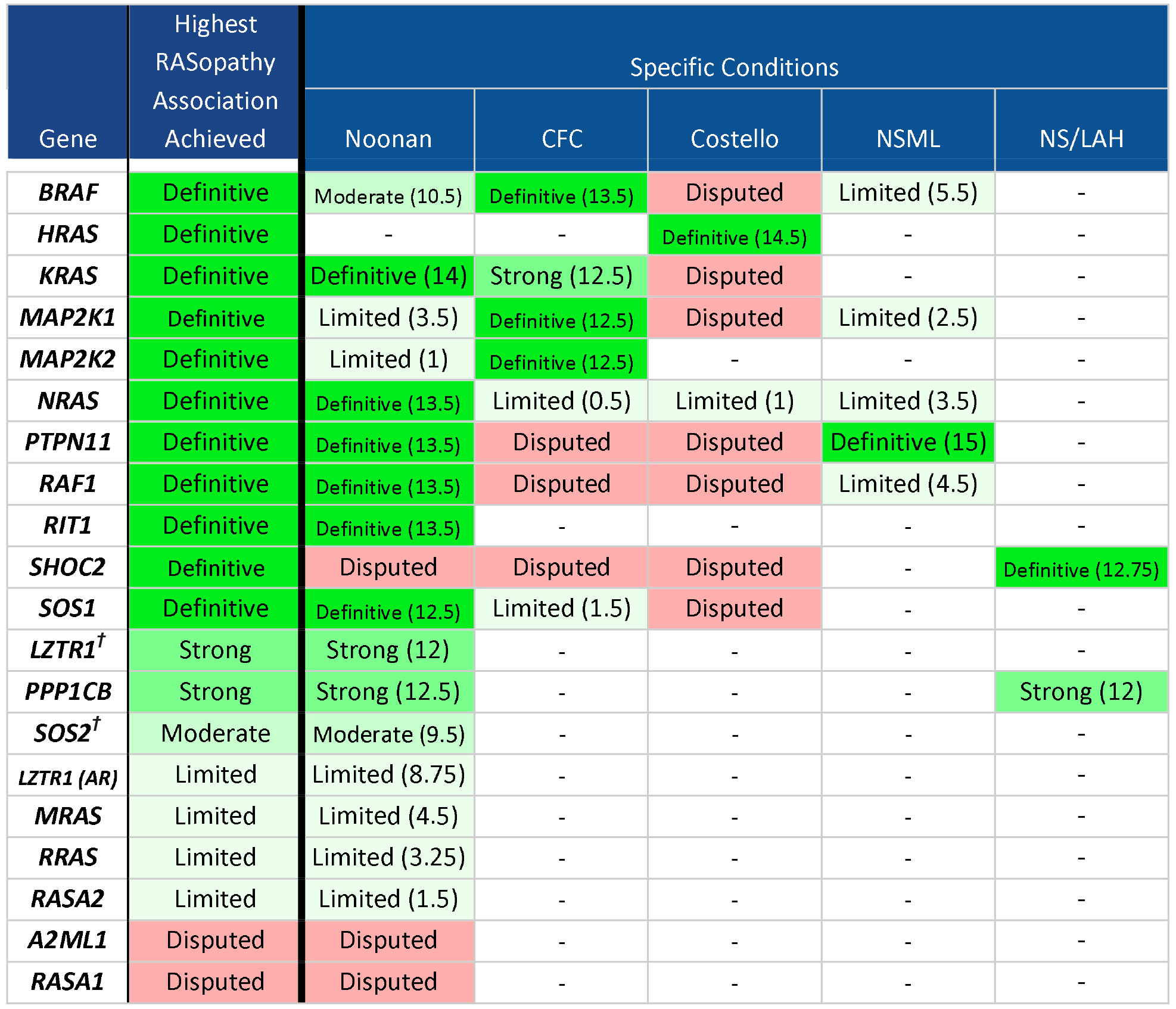
Classifications of each gene-disease association as curated by the ClinGen gene curation framework. Scored points for each association are shown in parentheses. “-”-No evidence, f-denotes a gene where publicly available data was used to supplement the published literature. AR - autosomal recessive inheritance.

| Gene | Highest RASopathy Association Achieved | Specific Conditions |  |  |  |  |
| --- | --- | --- | --- | --- | --- | --- |
|  |  | Noonan | CFC | Costello | NSML | NS/LAH |
| **BRAF** | Definitive | Moderate (10.5) | Definitive (13.5) | Disputed | Limited (5.5) | - |
| **HRAS** | Definitive | - | - | Definitive (14.5) | - | - |
| **KRAS** | Definitive | Definitive (14) | Strong (12.5) | Disputed | - | - |
| **MAP2K1** | Definitive | Limited (3.5) | Definitive (12.5) | Disputed | Limited (2.5) | - |
| **MAP2K2** | Definitive | Limited (1) | Definitive (12.5) | - | - | - |
| **NRAS** | Definitive | Definitive (13.5) | Limited (0.5) | Limited (1) | Limited (3.5) | - |
| **PTPN11** | Definitive | Definitive (13.5) | Disputed | Disputed | Definitive (15) | - |
| **RAF1** | Definitive | Definitive (13.5) | Disputed | Disputed | Limited (4.5) | - |
| **RIT1** | Definitive | Definitive (13.5) | - | - | - | - |
| **SHOC2** | Definitive | Disputed | Disputed | Disputed | - | Definitive (12.75) |
| **SOS1** | Definitive | Definitive (12.5) | Limited (1.5) | Disputed | - | - |
| **LZTR1<sup>†</sup>** | Strong | Strong (12) | - | - | - | - |
| **PPP1CB** | Strong | Strong (12.5) | - | - | - | Strong (12) |
| **SOS2<sup>†</sup>** | Moderate | Moderate (9.5) | - | - | - | - |
| **LZTR1 (AR)** | Limited | Limited (8.75) | - | - | - | - |
| **MRAS** | Limited | Limited (4.5) | - | - | - | - |
| **RRAS** | Limited | Limited (3.25) | - | - | - | - |
| **RASA2** | Limited | Limited (1.5) | - | - | - | - |
| **A2ML1** | Disputed | Disputed | - | - | - | - |
| **RASA1** | Disputed | Disputed | - | - | - | - |

## Discussion

### Gene Curation Workflow

While we utilized the ClinGen gene curation framework to assess these associations, there were some unique aspects of our process that should be noted. After primary curation, the scored literature evidence was presented to the ClinGen RAS EP for review. It included, for example, the number of probands with variants in the gene, number of *de novo* versus familial occurrences, functional data supporting the impact of the variants, and any phenotypic information available. Additionally, any publicly available variant data from diagnostic labs for genes that have been more recently associated with RASopathies, including *RRAS, LZTR1, RASA2, A2ML1, RASA1*, and *MRAS* was accessed via ClinVar (see **Supplementary Table S1).** The ClinGen RAS EP evaluated the evidence and commented on whether certain cases and experimental data should be scored. They also commented on the status of the current practices in the field and whether certain genes are traditionally thought of as being associated with certain phenotypes. In some cases, the ClinGen RAS EP asserted that certain associations may not align with current practices or may be inaccurate relative to the context of a particular case. For example, *BRAF* alterations are traditionally associated with CFC syndrome; however, *BRAF* alterations have also been identified in patients with phenotypes more typical of NS (Lee et al., 2011; Nystrom et al., 2008; Razzaque et al., 2007; Sarkozy et al., 2009; van Trier et al., 2016). In these instances, the RAS EP noted that the age of ascertainment, outdatedness of clinical assessments, and variable expressivity of the RASopathies should be considered when scoring these cases. After thorough discussion of the evidence, the RAS EP voted blindly on the classification of the clinical validity of the association. If a consensus was not achieved, the curation was discussed further until at least 80% of the RAS EP was in agreement. In these situations, specific language helped convey the reasons for the final decision of the RAS EP **(Supplementary Table S1).**

### Challenges of the Curation Process

The RASopathies are a complex group of disorders with substantial overlap in phenotypes. Additionally, the molecular mechanisms underlying the resultant RASopathy phenotype are not clearly correlated to specific variation within a gene. Our work highlights which discrete disorders have been shown to be associated with Ras/MAPK genes in the published literature. Unfortunately, a substantial amount of published literature does not provide enough information to differentiate between certain phenotypes such as NS versus CFC syndrome. This limitation is particularly related to the fact that most reported cases are children. Young age at presentation is a significant cause of disputable phenotypic classifications. Since the clinicians on the RAS EP could not clinically evaluate the patients described in publications themselves, we were reliant on the diagnosis provided in the literature.

Our decision to maintain separate associations for each gene and specific phenotype, instead of only providing a clinical validity classification for each gene’s association with a general RASopathy, was challenging. While there is an ongoing discussion in the field of the RASopathies as to whether these genes should be classified as a phenotypic spectrum under a broader disease entity, the clinical diagnoses currently provided to patients is historically supported as separate syndromes. Therefore, we felt it was important that our curations investigated each gene’s level of evidence for each of the five discrete syndromes. This allowed us to dispute weaker gene-disease associations that lacked convincing evidence. This process underscored the need for clear nosology guidelines for the RASopathies and the clinicians within the ClinGen RAS EP aimed to highlight specific phenotypic features that can assist in the differentiation of the RASopathies (see **Table 1).**

While we decided that it was important for each curation to assess the clinical validity of the specific phenotypes, we found that almost the entirety of experimental evidence currently in the literature does not provide differentiated support for a gene’s association with discrete phenotypes. For example, a mouse model with a knock-in *KRAS* p.Val14Ile variant which has been asserted to be in association with NS displayed short stature, cardiac abnormalities, craniofacial dysmorphisms, splenomegaly and myeloproliferative disorder, implicating the variant and gene’s causal role in development of a RASopathy phenotype (Hernandez-Porras et al., 2014). However, this mouse model does not provide evidence to discern between NS versus other RASopathies, especially regarding the known phenotypic overlap in humans. There are similar issues with studies showing that variants in these genes cause similar functional alterations in the Ras/MAPK pathway. For example, since the impact of *BRAF* variants associated with CFC syndrome and NS are both assessed by measuring increased phosphorylation of MEK or ERK, this evidence cannot be distinctly scored to support the association between *BRAF* and NS or CFC syndrome. While there are examples of individuals having the same variant and different clinical diagnoses (e.g. p.T241P, p.K499E, p.E501K), most evidence supports clinical variability is at least to some extent due to variant-specific differences in the phenotypic expression (e.g. BRAF mutations may be associated with more or less severe cognitive impairment) (Sarkozy et al., 2009). This genotype-specific variability of clinical expression likely results in the overlaps in gene:disease associations observed with the RASopathies. For example, an individual with a BRAF mutation and cognitive function in the normal or low-normal range is more likely to be classified clinically as NS.

Since there are several gene-disease associations that are considered uncommon in the field of RASopathies (*e.g.*, the associations of *KRAS* with Costello and *MAP2K1* with NS) many of the associations were disputed by our curations. However, if variants in the gene were associated with an “uncommon” disease but identified in a distinct risk profile within the phenotypic spectrum associated with this disease, the RAS EP typically deferred to the calculated classification of Limited or Moderate evidence, instead of disputing the association. In contrast, if an uncommon disease association was reported for a variant that had repeatedly been observed with the commonly-associated disease for its respective gene, the uncommon association was classified as Disputed. This applies, for example, to the associations of CFC syndrome with *PTPN11, SHOC2*, and *RAF1*.

### Examples of Challenging Curations

There were several curations in which the lack of clear gene:phenotype correlations merited careful review in the scoring of genetic evidence to support an association. Additionally, the identification of the same variant linked to multiple conditions revealed the lack of specificity in molecular evaluations to a given condition, yet still supported a general association to a RASopathy. Some examples are shown below but further details are provided in **Supplementary Table S1.**

#### *KRAS*-CFC syndrome

*KRAS* is traditionally associated with NS, but *KRAS* variants have also been identified in patients with a clinical diagnosis of CFC. Additionally, there is substantial phenotypic overlap between NS and CFC syndrome. For example, the cardiac abnormalities and facial anomalies possess a very similar spectrum (Table 1). While the severity of intellectual disability, frequency of severe ectodermal anomalies, and presence of neuropathy or other corticospinal tract anomalies can be used to differentiate NS from CFC syndrome, these distinguishing features can still be observed in NS and are age-dependent. Therefore, despite sufficient genetic evidence and repetition over time for *KRAS* and CFC syndrome to establish a Definitive classification, these complexities led clinicians in the RAS EP to caution against classifying CFC syndrome as Definitively associated with *KRAS*. The RAS EP chose to classify the association as Strong.

With the evidence provided in the literature, it was not always convincing to the RAS EP that a case described as CFC syndrome with a *KRAS* variant would not develop features more consistent with NS over time due to the fact that many of the patients were less than 10 years of age (Adachi, Abe, Aoki, & Matsubara, 2012; Niihori et al., 2006; Zenker et al., 2007). Of note, there were cases in the literature who were as old as 26 presenting with features more consistent with CFC syndrome (Sovik et al., 2007). This scenario exemplifies that a molecular diagnosis should not dictate the final clinical diagnosis, but encourage clinicians to scrutinize the phenotypic presentation of the individual over time. While our consistent approach was to accept the diagnoses within the peer-reviewed literature provided and score cases if the variants themselves had sufficient evidence for pathogenicity, in this particular case, the RAS EP decided that the uncertainty surrounding the cases provided exceptional reasons to downgrade the classification from Definitive to Strong. We cite variant overlap between CFC syndrome and NS patients, variable expressivity, outdated assessments, and young age of ascertainment of diagnoses of affected individuals as reasons for downgrading this classification from Definitive to Strong **(Supplementary Table S1).**

#### *LZTR1* Associations

*LZTR1* is a gene that has been more recently (< 3 years from first publication) associated with the RASopathies. The published cases to date have presented with a phenotype consistent with NS; however, the mechanism of inheritance is variable. Autosomal dominant (AD) variants are exclusively missense (Chen et al., 2014; Yamamoto et al., 2015), while the autosomal recessive (AR) variants are predominantly predicted loss of function (LOF) with at least one hypomorphic allele being present in patients with biallelic mutations (Johnston et al., 2018). It could be inferred that biallelic LOF or a GOF, potentially as dominant-negative mechanism, missense variant can lead to the Noonan phenotype. Of note, *LZTR1* is not LOF constrained according to ExAC data, suggesting that these variants may be tolerated (exac.broadinstitute.org)(Lek et al., 2016). Further identification and publication of *LZTR1* RASopathy cases are necessary to further confirm these disease mechanisms. Given that variants in *LZTR1* have been shown to segregate with an NS phenotype in both inheritance patterns and apparently due to different disease mechanisms, we chose to split the curations into scores for both the AR and AD associations of *LZTR1* with the RASopathies. Furthermore, since there have only been three publications to date, we utilized publicly available, yet not formally published data from diagnostic laboratories in ClinVar to achieve the most up-to-date evidence score that reflects the current acceptable association in the field. Evidence of multiple confirmed *de novo* occurrences primarily justified the score of 12 for genetic evidence. Lack of experimental evidence and replication overtime prevents a Definitive association, thus the AD association is currently assessed as Strong, but is expected to move to Definitive in the near future if no contradictory evidence emerges.

#### *BRAF/MAP2K1/MAP2K2*: Noonan syndrome

Variants in these genes have repeatedly been associated with NS, but in many publications, the rationale and basis for the clinical diagnosis of NS over CFC is unclear. As mentioned previously, many of the variants associated with NS in these genes have also been reported in association with CFC. This may be further complicated by the lack of consensus criteria and nosology for differentiating NS from CFC syndrome clinically. Moreover, functional differences between NS- and CFC syndrome-associated *BRAF/MAP2K1/MAP2K2* variants have not been established, so far. These reasons may support categorizing and classifying these genes with a broader disease entity that encompasses both NS and CFC (i.e. a RASopathy), but the RAS EP felt it was necessary to maintain the traditional disease conditions and assess the strength of evidence for each phenotype associated with these genes in the literature. It should be noted that none of the associations of these genes with NS reached the maximum score for genetic evidence, thus indicating further evaluation of evidence is needed in the future. The RAS EP noted that some variants in *BRAF* and *MAP2K1* were consistently associated with NS alone while other variants had broader assertions, indicating the potential for specific genotype:phenotype correlations to be elucidated over time with further experimentation and exploration through functional assays. The RAS EP recognized that the NS diagnosis in individuals carrying variants in genes typically associated with CFC syndrome is a major point of discussion when attempting to classify gene-disease associations among RASopathies. The RAS EP asserted that a phenotypic diagnosis based exclusively on the molecular diagnosis from a gene (and not genotype) level in this area would not appropriately reflect the broad phenotypic spectrum associated with mutations in *BRAF, MAP2K1* and *MAP2K2.*

#### *PTPN11/BRAF/RAF1*: NSML

There is a Definitive association for *PTPN11* variants with NSML. Unlike other associations, there is robust evidence for the major NSML-associated *PTPN11* variants (p.Y279C and p.T468M) displaying clearly different functional properties (reduced phosphatase activity) compared to NS-associated variants. However, this has not been shown for all NSML-associated *PTPN11* variants (Fragale et al., 2004; Kontaridis et al., 2006; Noda et al., 2016; Oishi et al., 2006; Oishi et al., 2009; Yu et al., 2014). Moreover, young individuals lack clear distinguishing phenotypic characteristics between NS and NSML so precise diagnoses may be difficult, thus explaining the reported associations of NS with “classical” NSML-associated variants in the literature. In addition to the distinct functional impact of NSML-associated *PTPN11* variants, there is also a risk profile that distinguishes NSML from NS (*e.g.* high risk of HCM, risk of hearing deficits). The differentiation is less clear for *BRAF* and RvAFl-associations with NSML, and no functional impact of NSML-associated *BRAF/RAF1* variants has been established in these genes. Furthermore, the RAS EP asserted from their own experience that even the skin phenotype associated with *BRAF* and *RAF1* is distinct from the multiple lentigines phenotype seen in patients with “classical” NSML-related *PTPN11* variants.

#### *SHOC2* with NS/CFC syndrome/CS

Primary curation showed “Limited” evidence for an association of *SHOC2* with NS, CFC syndrome and CS. After discussion, the RAS EP decided to classify all other disease associations for *SHOC2* except NS/LAH as Disputed given the primary common pathogenic variant (p.Ser2Gly) in this gene has been reported over 100 times with a clinical diagnosis of NS/LAH. In addition, this mutation has distinct functional consequences, and the resemblance of the phenotype associated with variants in *PPP1CB*, a known direct interaction partner with *SHOC2*, further supports that NS/LAH is a discrete entity specifically related to alterations in this signaling complex module (Aoki, Niihori, Inoue, & Matsubara, 2016; Rauen, 2013). While it is true that young patients with NS/LAH may have a phenotype reminiscent of CS or CFC syndrome (*e.g.*, hair abnormalities, severe feeding issues, and more severe developmental issues than usually seen in NS), the phenotype evolves to become more Noonan-like in older children. Considering the specific phenotype of NS/LAH, the RAS EP judged that there is insufficient evidence for classifying the diagnosis as CFC syndrome, CS or NS instead of NS/LAH in any of the published patients. The RAS EP concluded that the reason for divergent associations in the literature is related to the fact that authors were likely only considering the categories NS, CFC syndrome, and CS before NS/LAH existed as a separate clinical entity. Therefore, all the cases that have been presented to date, should be considered under the NS/LAH phenotype, thus disputing the association of *SHOC2* with NS, CFC syndrome, and CS.

## Conclusion

Given the complexities surrounding the RASopathies, we believe that our curation processes thorough assessment of the current published and publicly available evidence in ClinVar provides insight into the current issues in the field. By utilizing the ClinGen gene curation framework and expert review from members of the ClinGen RAS EP, we compiled evidence and commentary on almost 100 RASopathy gene-disease associations. In choosing to assess the evidence for each specific phenotype for each gene, we provide a comprehensive review of several controversial associations that we believe will be useful in both clinical and molecular diagnoses. Additionally, the difficulties in validating gene-disease associations among RASopathies demonstrate that all authors that publish on genotype-phenotype associations in RASopathies should rigorously substantiate rare phenotype associations with detailed and comprehensive clinical assessments for the purported diagnosis. While more efforts are needed to address clear nosology of these disorders to aid in clinical diagnosis, the possibility of a diagnosis of a Definitively-associated phenotype with a given gene (per the classifications established here) should be extensively scrutinized and essentially ruled out with valid phenotypic evidence before asserting a disease association with only Limited or Moderate classification.

While we aimed to provide the community with the most up-to-date associations, we caution that these associations are just a snapshot of the evidence currently available. For example, the RAS EP predicts that the non-definitive associations of newly recognized genes like SOS2, PPP1CB, LZTR1, and MRAS are likely to become definitive overtime as sufficient evidence accumulates in the literature. On the other hand, limited associations for genes historically studied are likely to remain limited or even potentially disputed on re-evaluation. As additional evidence becomes available for expert review, the RAS EP will aim to update and refine classifications on the Clinical Genome Gene-Disease Validity website (https://www.clinicalgenome.org/curation-activities/gene-disease-validity/results/) and we encourage readers to publish any case-level or experimental evidence that may support, challenge, or inform this curation project and its clinical validity classifications.

## Acknowledgments

The authors would like to thank Heidi Rehm, Jonathan Berg, Laura Milko, Courtney Thaxton, Erin Riggs, Marina DiStefano and the additional members of the ClinGen Gene Curation Working Group and Clinical Domain Oversight Committee for their points of feedback on the gene curation approach and manuscript.

## Authorship Contributions

Gene validity curation was performed by BJC and ARG and overseen by LMV and MZ. All authors reviewed and approved clinical validity classifications. HC, BDG, KWG, KAR, and MZ provided clinical expertise and differentials for each RASopathy condition. Finalization of clinical validity classifications and manuscript content was overseen by LMV and MZ.

